# Design of a comprehensive microfluidic and microscopic toolbox for the ultra-wide spatio-temporal study of plant protoplasts development and physiology

**DOI:** 10.1101/526889

**Authors:** K. Sakai, F. Charlot, T. Le Saux, S. Bonhomme, F. Nogué, J.C. Palauqui, J. Fattaccioli

## Abstract

**Background:** One of the main features of plant cells is their strong plasticity, and their propensity to regenerate an organism from a single cell. Plant protoplasts are basic plant cells units in which the pecto-cellulosic cell wall has been removed, but the plasma membrane is intact. One of the main features of plant cells is their strong plasticity, which in some species, can be very close from what is defined as cell totipotency. Methods and differentiation protocols used in plant physiology and plant biology usually involve macroscopic vessels and containers that make difficult, for example, to follow the fate of the same protoplast all along its full development cycle, but also to perform continuous studies of the influence of various gradients in this context. These limits have hampered the precise study of regeneration processes.

**Results:** Herein, we present the design of a comprehensive, physiologically relevant, easy-to-use and low-cost microfluidic and microscopic setup for the monitoring of *Physcomitrella patens* (*P. patens*) growth and development on a long-term basis. The experimental solution we developed is made of two parts (i) a microfluidic chip composed of a single layer of about a hundred flow-through microfluidic traps for the immobilization of protoplasts, and (ii) a low-cost, light-controlled, custom-made microscope allowing the continuous recording of the moss development in physiological conditions.

We validated the experimental setup with three proofs of concepts: (i) the kinetic monitoring of first division steps and cell wall regeneration, (ii) the influence of the photoperiod on growth of the protonemata, and (iii) finally the induction of leafy buds using a phytohormone, cytokinin.

**Conclusions:** We developed the design of a comprehensive, physiologically relevant, easy-to-use and low-cost experimental setup for the study of *P. patens* development in a microfluidic environment. This setup allows imaging of *P. patens* development at high resolution and over long time periods.

## BACKGROUND

Protoplasts are basic plant cells units (De Smet) in which the pecto-cellulosic cell wall has been removed, but the plasma membrane is intact. These fragile entities can be isolated from various parts of the adult plant (leaves, etc.) using enzymatic treatments (pectinase, cellulase) (Wiszniewska, 2012). The softness of the shallow protoplast membrane allows making genetic manipulation following the direct insertion of various molecules (DNA, etc.), objects (droplets, etc.), organelles (chloroplasts, mitochondria) or nuclei within the cell (De Smet; Reski, 1998). In proper culture conditions, protoplasts can dedifferentiate, divide, re-differentiate in various cell types, to finally lead to the regeneration of a full organism. Protoplasts are hence interesting models both in the perspective of single-cell and developmental studies. However, culture methods and differentiation protocols used for plant physiology and plant biology usually involve macroscopic vessels and containers (Cove et al., 2009a) that make difficult to follow the fate of the same protoplast all along its full development cycle in a continuous manner.

Thanks to their ability to recreate *in-vitro* environments at the cellular level that mimic a tissue, and also observing in real-time subcellular and cellular modifications that take place during growth or development, microfluidic devices have been used as monitoring and culturing tools for insect embryos(Levario et al., 2013), mammalian (Mehling and Tay, 2014), or bacterial cells (Eland et al., 2016). In plant science however, few microdevices have been developed so far (Sanati Nezhad, 2014). Among them, the vast majority of the literature deals with root or pollen tube immobilization and studies (Bérut et al., 2018; Gooh et al., 2015; Grossmann et al., 2011; Kamimura et al., 2018; Massalha et al., 2017; Meier et al., 2010; Park et al., 2014; Park et al., 2017), and half a dozen only address the question of the manipulation and observation of protoplasts (Bascom et al., 2016; Grasso and Lintilhac, 2016; Ju et al., 2006; Ko et al., 2006; Schaap et al., 2016; Wu et al., 2011; Zaban et al., 2014).

Among them, Ko *et al.* (Ko et al., 2006) developed a microchip for the trapping, the culture and the fusion of tobacco protoplasts. Trapping takes place in a series of PDMS posts acting as a filter and stopping protoplasts within a microchannel. In addition, Zaban *et al.* (Zaban et al., 2014) developed a microfluidic system to impose chemical cues to regenerating protoplasts entrapped individually in an array of small PDMS wells, pinpointing the importance of the auxin gradient on the directionality of the cell wall regeneration. While these two studies were focused on the regeneration of protoplasts on short timescales, Bascom *et al.* (Bascom et al., 2016) presented recently a microfluidic setup allowing the long-term culture of moss protoplasts in static culturing condition, i.e. in absence of flow inside the chamber. While controlled light conditions, such as photoperiodicity or energy flux, are a stringent constraint for the physiological development of protoplasts (Jenkins and Cove, 1983) in a laboratory environment, none of the solutions published so far provides satisfactory lightning solutions during the microscopic observations, probably because conventional microscopes are not easily usable in plant-compatible illumination environmental setups. Understanding the cell fate during plant development thus necessitates the design of novel monitoring and observation techniques to address these questions.

Herein, we present the design of a comprehensive, physiologically relevant, easy-to-use and low-cost microfluidic and microscopic setup that takes into account all the above constraints related to plant growth and development studies and overcomes current experiment limitations. For the sake of demonstration, we used a simple plant model, the moss *Physcomitrella paten*s, which has the ability to easily regenerate individuals from isolated protoplasts (Cove et al., 2009a; Cove et al., 2009b). The experimental solution we developed is made of two parts : (i) a microfluidic chip composed of a single layer of about a hundred flow-through microfluidic traps (Karimi et al., 2013) that immobilize protoplasts without hindering their development in a sterile and portable environment; (ii) a microscope inserted in a light-controlled incubator that allows a controlled illumination and the continuous recording of the development over time. After having given a detailed description of experimental setup and the design rationales, we validated it on three proofs of concepts characterized by different spatiotemporal scales: the kinetic monitoring of first division steps and cell wall regeneration, the influence of the photoperiod on growth of the protonema, and finally the induction of leafy buds using a cytokinin as a well described phytohormone.

## RESULTS

### Size distribution of *P. patens* protoplasts and spores

Starting from 1-week-old *P. patens* protonema, protoplasts are isolated by enzymatic digestion with driselase, using protocols detailed in the **Materials and Method** section. After digestion of the cell wall, protoplasts are very fragile and are hence suspended in an iso-osmotic 8.5 wt% mannitol solution. To remove undigested protonema and cellular debris, the suspension is filtered using a set of sieves of decreasing sizes (80 µm, 40 µm). At the end of the isolation process, a representative picture of the suspension is shown in **Figure 1A**. By microscopy and image analysis, we measured the size distribution of a typical protoplast suspension, shown in **Figure 1C**. Protoplasts have a homogeneous size distribution characterized by an average size equal to 32 ± 5 µm. In parallel to protoplast isolation, we also measured the size distribution of *P. patens* spores shown in **Figure 1B**. As compared to protoplasts, **Figure 1C** shows that spores have the same average diameter but are more homogeneous in size.

**Figure 1:**
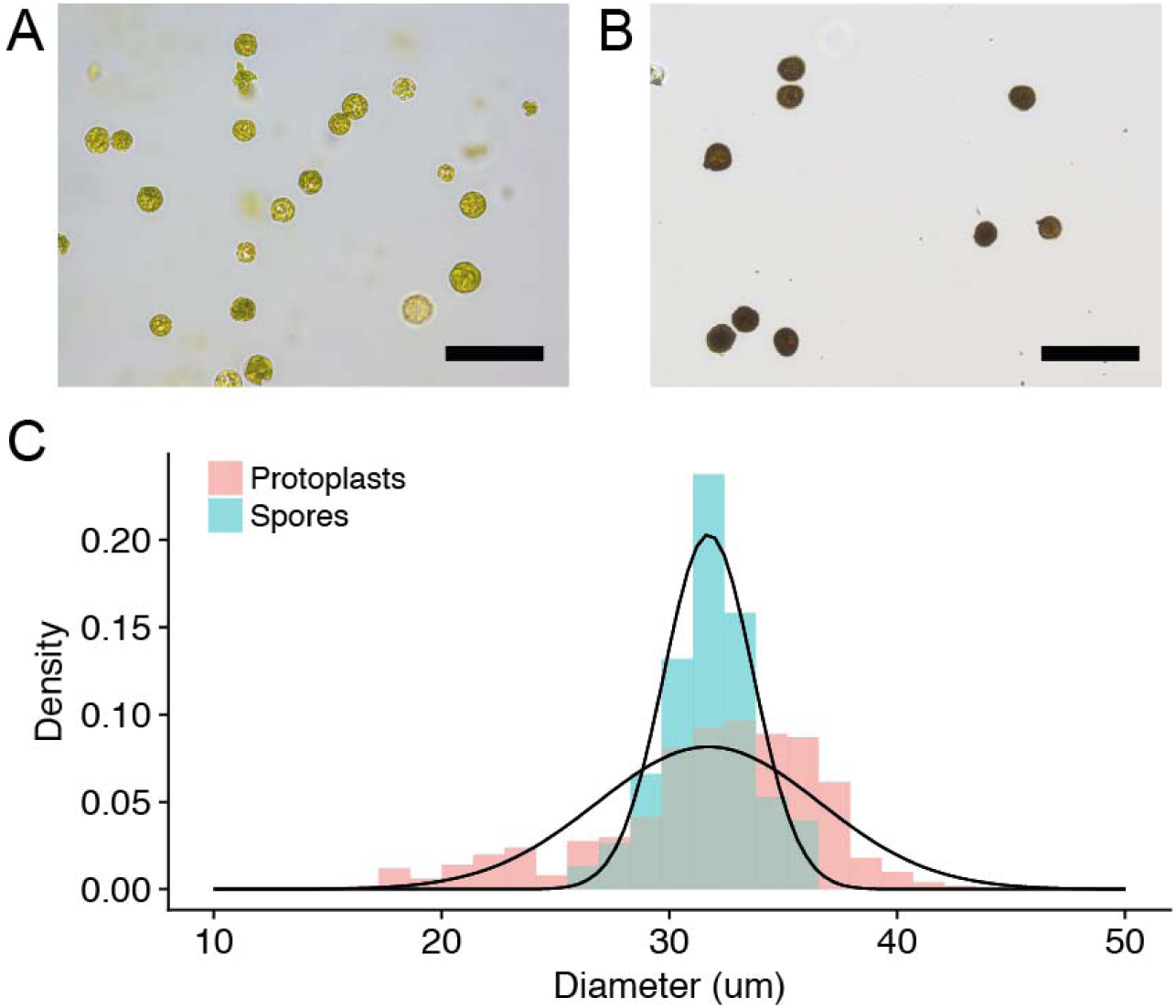
Representative brightfield microscopy pictures of a suspension of *P. patens* protoplasts (A) after digestion and purification, and (B) spores (C) Size distributions of protoplasts (Red) and spores (Blue) suspension. A gaussian fit of the size distributions gives a diameter of 32 ± 5 µm and 32 ± 2 µm respectively. Scalebars : (A) 100 µm, (B) 100 µm

### Hydrodynamic traps design and microfluidic setup

The chip is made of a 2D microfluidic chamber containing a regular array of 14 staggered lines of 8 hydrodynamic flow-through traps (**Figure 2A**). The height of the microfluidics chamber is set to 45 µm. Microchamber height and trap dimensions have been chosen in accordance with the size distribution of the protoplasts, with the general compromise to optimize immobilization and mechanical stability while avoiding a too strong confinement that could make loading difficult and eventually alter protoplast development. Traps are U-shaped, and after several optimization steps, we designed them with a backside and two lateral opening to allow the culture medium flowing easily through the tiny structures, thus avoiding the existence of dead volumes; and to give freedom to the developing protoplasts to grow.

**Figure 2:**
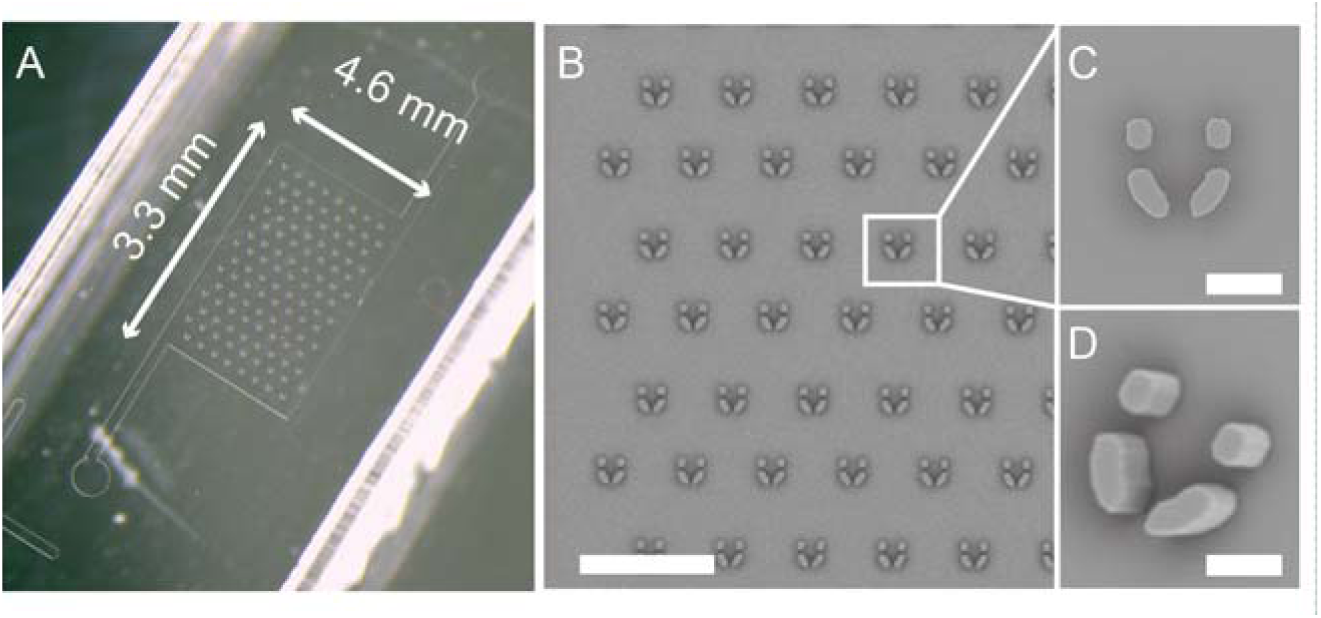
(A) Complete binocular view of the microchip. Length = 4.6 mm, width = 3.3 mm (B) Enlarged SEM view of the chamber containing the trap array. The vertical and horizontal spacings of the traps are respectively 200 µm each. Scalebar = 500 µm. (C) Top and (D) tilted SEM view of a trap. The traps are U-shaped (inner length = 60 µm, inner width = 45 µm) and have 2 lateral and one backside (15 µm) opening to allow the medium to flow through the structure. The chamber height is set to 45 µm. Scalebar = 40 µm.

Traps shown in **Figure 2** have an inner length of 60 µm, an inner width of 45 µm, a wall thickness of 20 µm, and opening widths equal to 15 µm **(Figure 2B-C**). Microfluidic trap arrays are fabricated using soft-lithography techniques (Xia and Whitesides, 1998) detailed in the **Materials and Methods** section. After PDMS molding and creation of openings for tubing insertion, PDMS chips are bonded to glass-bottom Petri dishes, as shown in **Figure 2C**, to ease manipulation and microscopic observation. To get a stable hydrophilic coating of the inner wall of the microchip and avoid trapping of air bubbles, microfluidic chambers are rinsed with an hydrophilic block-polymeric surfactant solutions (0.2 wt% of Pluronic F68 in water) prior to the experiments.

### Protoplast loading and immobilization

After membrane digestion and sieving, protoplasts are first diluted at the concentration of 10^5^-10^6^ mL^-1^ in an iso-osmotic buffer made from PpNH4, 6.6 wt% of glucose and 0.5 wt% of mannitol. Then, they are inserted in a microtube connected to a computer-controlled pressure regulator. Protoplasts are injected in the microchamber using a pressure drop of 10-15 mbar during 2 to 3 minutes, low enough to avoid a strong shear rate of the protoplasts.

**Figure 3A** shows that each individual trap contains 1 to 5 protoplasts. Trap loading does not strictly follows a Poisson distribution since the presence of an immobilized protoplast in a trap modifies the flow-through hydrodynamic stream, hence decreasing the probability to trap a second cell (Di Carlo et al., 2006). **Figure 3B** shows that for the same set of experiments, the filling rate, measured by counting the number of traps containing at least one protoplast, is close to 90%. Starting from these values, the trapping efficiency can be easily tuned by playing with the cell density and the time of loading.

**Figure 3:**
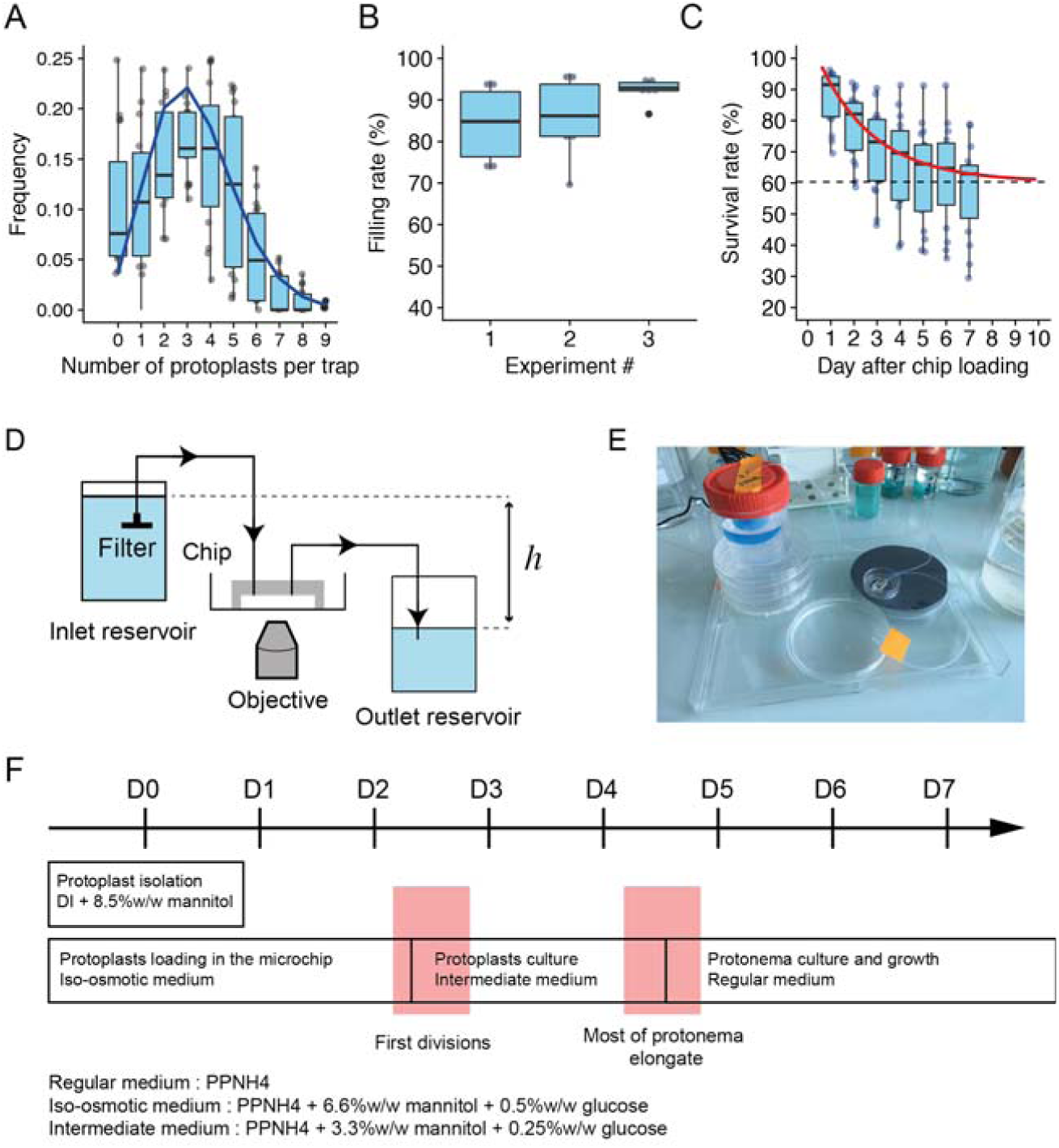
(A) Histogram of the number of protoplasts per trap (pooled data, n=6, 3 replicates). The blue line corresponds to a Poisson distribution with an average number λ=3.3. (B) Comparative dotplots of the filling rates of the microfluidic chips, for three sets of independent experiments. The filling rate is defined as the percentage of traps for a given device containing at least one protoplast after the completion of the injection step. The filling rate is equal 87,4 ± 8,4 protoplasts per trap (pooled data, n=6 devices, 3 replicates). (C) Evolution of the protoplasts survival rate in a microchip as a function of time. Day 1 corresponds to 1 O/N after protoplast loading. During the first 4 days of culture in the chip, the survival rate decreases, and it stabilizes from day 7 at 62 % in average (pooled data, n=5, 3 replicates). The black line corresponds to an exponential fit performed on the mean values of the survival rate. (D) Schematic representation of the gravity-driven fluid control setup and its connection to the microchip. The medium flow rate is set by the hydrostatic pressure difference between both reservoirs. To avoid debris and salt crystals going into the chamber, the incoming tubing entrance is connected to a 0.2µm filter. (E) Overall view of the experimental setup. (F) Timeline of the culture media changes in the microchip during the first week after cell wall digestion and isolation from protonema (Day 1).

### Regeneration, growth and differentiation procedure

After protoplast loading in the microchip, the fluidic control system is exchanged for a simpler and more transportable setup, based on a hydrostatic pressure (Komeya et al., 2017) difference between two reservoirs [27], as shown on the schematic view from **Figure 3D** and the **Figure 3E** photograph. The PDMS-on-glass chambers inlet is connected to an upstream source of medium reservoir, and the outlet is connected to a downstream waste disposal reservoir. Culture media are kept in large (100 mL) and sterile plastic containers that are used for several days in a continuous manner without external energy supply. After careful screening of some of the tubing available on the market, we chose to connect reservoirs and the chip using tubes made from a fluorinated polymer (PTFE), as they give best results in terms of contamination and bubble avoidance used silicone tubing (e.g. Tygon). The height difference between the culture media menisci in the inlet and outlet reservoir is set to 4 cm, which corresponds to a flow rate of 0.36 mL.hr^-1^ (6 µL.min^-1^)

Salts such as tartrates or mannitol both present at high concentration, both in PpNH4 medium at some stages of the experiment, tend to nucleate crystals (Kelly et al., 1961) from the walls of the microchip. These crystals modify the hydrodynamics resistance of the chip, degrade the microscopic image quality and can ultimately impede culture media flowing in through the chamber. We solved this issue by placing a 0.2 µm sterile filter connected to the inlet tubing and plunging into the inlet reservoir, as shown in **Figure 2 : D.**

### Survival rate of protoplasts in the microdevice

To allow a reproducible cell wall regeneration and then protoplasts division and growth, we optimized the culture media exchange routine, as shown in **Figure 3F**. During the loading step that takes place in the microchip and until the occurrence of the first cell divisions, protoplasts are maintained in an iso-osmotic medium made from PpNH4 medium supplemented with 6.6 wt% of mannitol and 0.5 wt% of glucose. Then, culture medium is replaced for ca. 2 days by a solution of PpNH4medium containing intermediate half concentrations of 3.3 wt% of mannitol and 0.25 wt% of glucose. Finally, when most of the protonema have started their elongation process, the culture medium is definitely replaced by pure PpNH4 medium.

Following this routine and during the first week after loading and immobilization of protoplasts in the chip, we quantified their survival rate by manually identifying and counting the proportions of traps containing at least one living protoplast. The status of the protoplasts is assessed by regularly monitoring the integrity of their membrane and chloroplasts, and their ability to develop over time. Survival rate 2 days after chip loading is of the order of 80 %, and stabilizes around 60 % after 5 days of culture in the microfluidic device, as shown by the histogram in **Figure 3E**

### First divisions and cell wall regeneration

Reconstitution of the pecto-cellulosic cell wall of *P. patens* protoplasts usually takes place during the first days after isolation and regeneration (Stumm et al., 1975). During that time, protoplasts are very fragile, which makes usually difficult the observation of the first steps of the development. Using the discrete array of traps in the microfluidic setup, we are able to immobilize the protoplasts and keep them in a gentle flow of iso-osmotic medium without degradation of the observations conditions. **Figure 4A** shows a laser-scanning confocal microscopy image of a trap containing 4 H2B-mRFP / microtubules-GFP protoplasts and observed for 48 hours after immobilization. **Figure 4B** shows a time-lapse recording of the first cytokinesis event of one of the protoplasts at a 20 min temporal resolution. We see the formation of the mitotic spindle and nuclei division, thanks to the histone labeling (in purple). Using calcofluor-white, a non-specific fluorescent stain that binds strongly to structures containing cellulose (Monheit et al., 1984), we followed the cell wall reconstitution as a function of time. **Figure 4C** shows that for each protoplast, the intensity of the cell wall is spatially heterogeneous and increases over time. This observation is in accordance to former reports from the literature (Jenkins and Cove, 1983).The average integrated intensity, shown in **Figure 4D**, increases linearly with time, while the coefficient of variation (CV), defined as the ratio of the standard deviation to the mean, saturates on long time scales, thus indicating an spatial homogenization of the cell wall after ca. 10 hours of experiment. **Figure S 2** shows that while protoplasts having experienced a long observation under a confocal microscope are unable to grow, protoplasts trapped in adjacent traps are growing normally with time.

**Figure 4:**
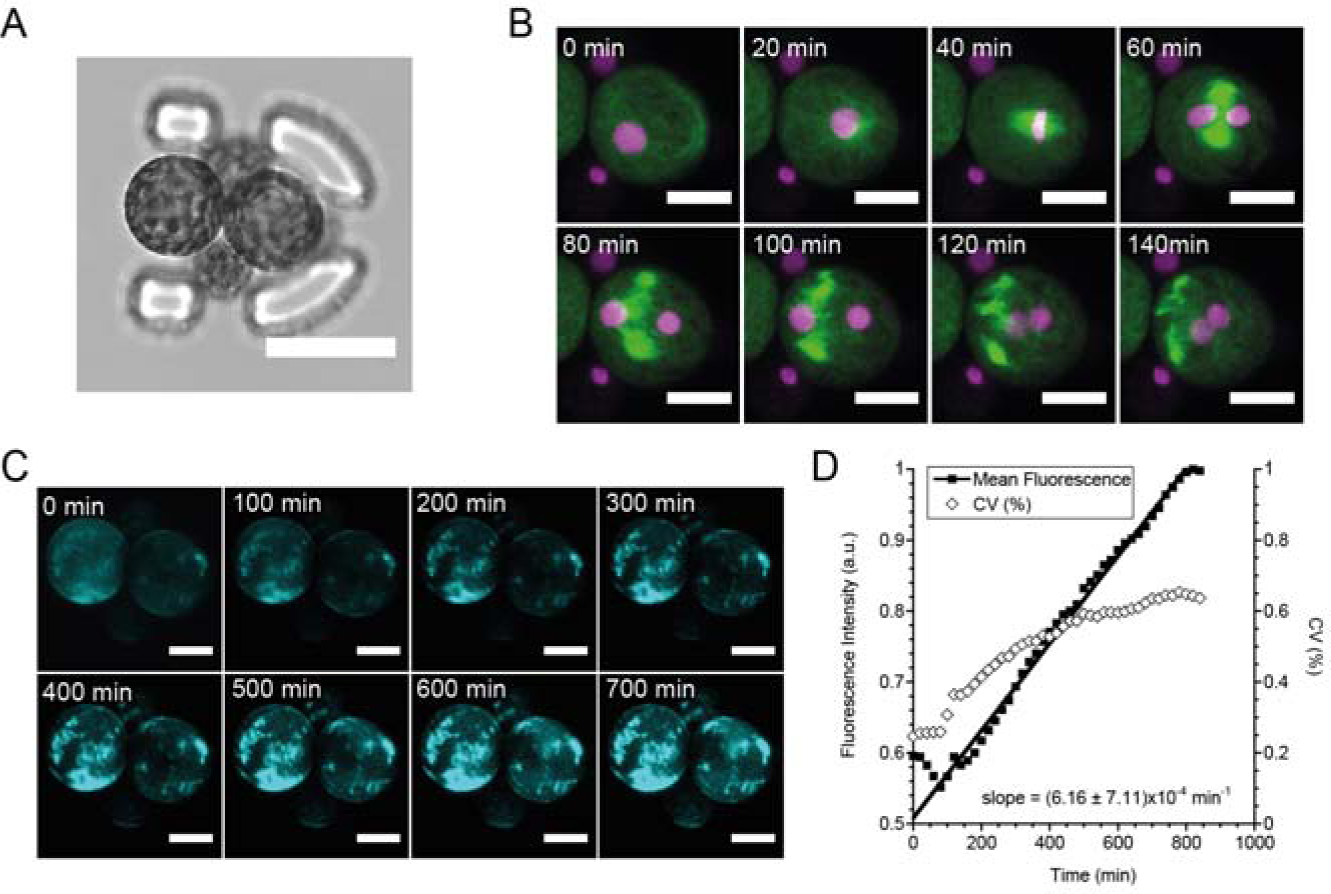
(A) Confocal image (transmission mode) of a trap containing four H2B-mRFP / microtubules-GFP protoplasts. Scalebar = 40 µm. (B) Confocal time-lapse imaging of the first division event of the protoplast #2 (purple : H2B-mRFP, green : microtubules-GFP). The GFP channel is in maximum projection mode. Scalebar : 20 µm. (C) Confocal time-lapse imaging of the cell wall reconstitution around the two protoplasts, using calcofluor-white as a dye. Representation is shown in maximum projection mode. Scalebar : 20 µm. (D) Mean fluorescence intensity of protoplast 1 over time in the calcofluor-white channel. CV is defined as the ratio of the standard deviation to the mean. The mean intensity increases linearly with time.

### In-house microscope design

To allow long-term observation of *P. patens* development in the microsystem, we have built a custom-made microscope, small enough to be inserted in a simple day/night incubator. A schematic view of the instrument design is shown in **Figure 5A**, and picture of the microscope is shown in **Figure 5B**. Inside the incubator, made from a commercial black plastic storage box, the night/day illumination is created by a LED tile controlled by a mechanical time switch. The microscope itself is made from common optical mounting elements, a CMOS color camera for imaging, a xy manual translation stage and an Arduino-controlled LED for the intermittent illumination of the sample all along the experiment. The LED element is built by the manufacturer with a collecting lens of large numerical aperture, ca. 60°, that makes illumination homogeneous in the field of view without the need to use a conventional condenser. To immobilize the glass-bottom Petri dish containing the microchip, we designed a 3D-printed adaptor for the xy-stage. A temperature sensor is also connected to the Arduino board for the continuous recording of this parameter during the experiment. The flowchart of the recording routine is shown in **Figure 5C**. To ease the experiments, we developed a graphical user interface (Matlab) for the control of the LED intensity, the image timelapse parameters, and the recording of the pictures taken with the camera. Currently, the microscope is built with a 10x plan-corrected objective with a low numerical aperture and without any tube lens. The optical setup has a field of view sufficient to observe 1 to 3 developing plants. Our setup can work continuously for several weeks for a given sample without noticing any defocusing, nor any detrimental effect of the presence of the LED element on the plant growth and development. **Figure 5D** shows a time-lapse recording of the development of chloronema for 40 hours in photoperiodic conditions, where tip growth and side-branching are clearly visible despite the simplicity of the setup.

**Figure 5:**
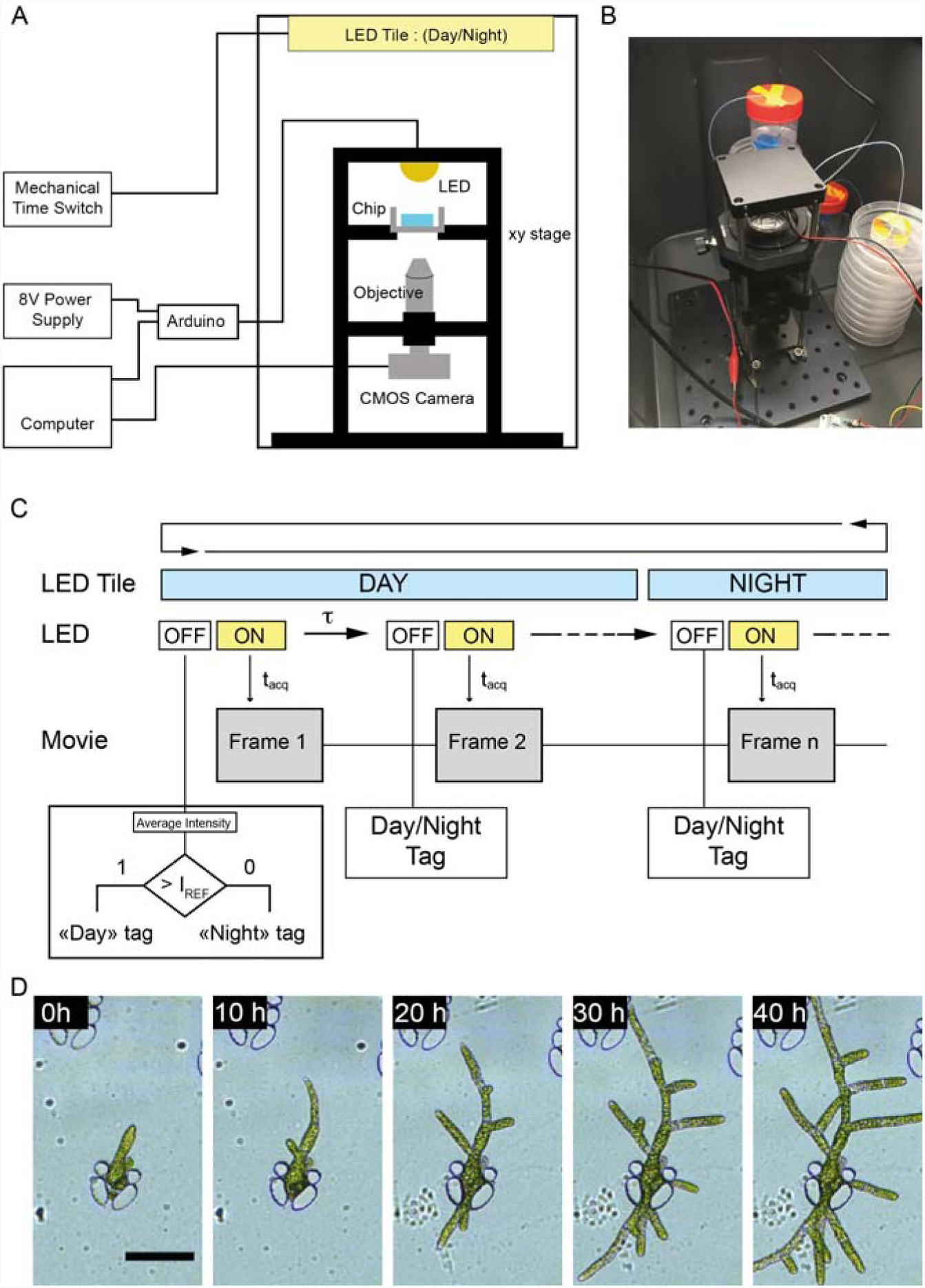
(A) Schematic representation of the in-house microscope built for long-term studies. The white illumination is provided by a white LED controlled with an Arduino micro-controller. The LED is turned on intermittently for a short amount of time to allow the picture recording by the CMOS camera. The camera is controlled by a Matlab script and the Micro-Manager software. The microscopic setup is encased in a box equipped with a LED tile on the ceiling, used to set the experimental photoperiod (night : 8h, day : 16h). The chamber temperature is recorded all along the experiment using a thermocouple sensing module. (B) Picture of the experimental setup during an experiment. (C) Flowchart of the movie recording routine and frame tagging with the photoperiod name. (D) Time-lapse microscopic recording of the growth of *P. patens* in the microdevice achieved with the custom-built setup, from day 6 after immobilization. Scalebar = 100 µm.

### Influence of the light cycle on protonemal growth

Using the custom-made microscope, we monitored the influence of the photoperiod on the protonemal growth of immobilized protoplasts. 7 days after isolation and culture in the microsystem, microfluidic samples are inserted in the microscope and observed continuously for ca. 80h with a temporal resolution of 10 minutes, either under a 16 h light/8 h darkness photoperiod or under continuous illumination. Under a photoperiodic culture condition, **Figure 6A** shows that the absence of light strongly decreases the tip elongation process, whereas growth restarts after a short lag-time when illumination takes place. Under continuous illumination conditions, growth is globally constant, as shown in **Figure 6B**. Growth rate in both conditions, measured during growing period, is of the order or 3µm.h^-1^.

**Figure 6:**
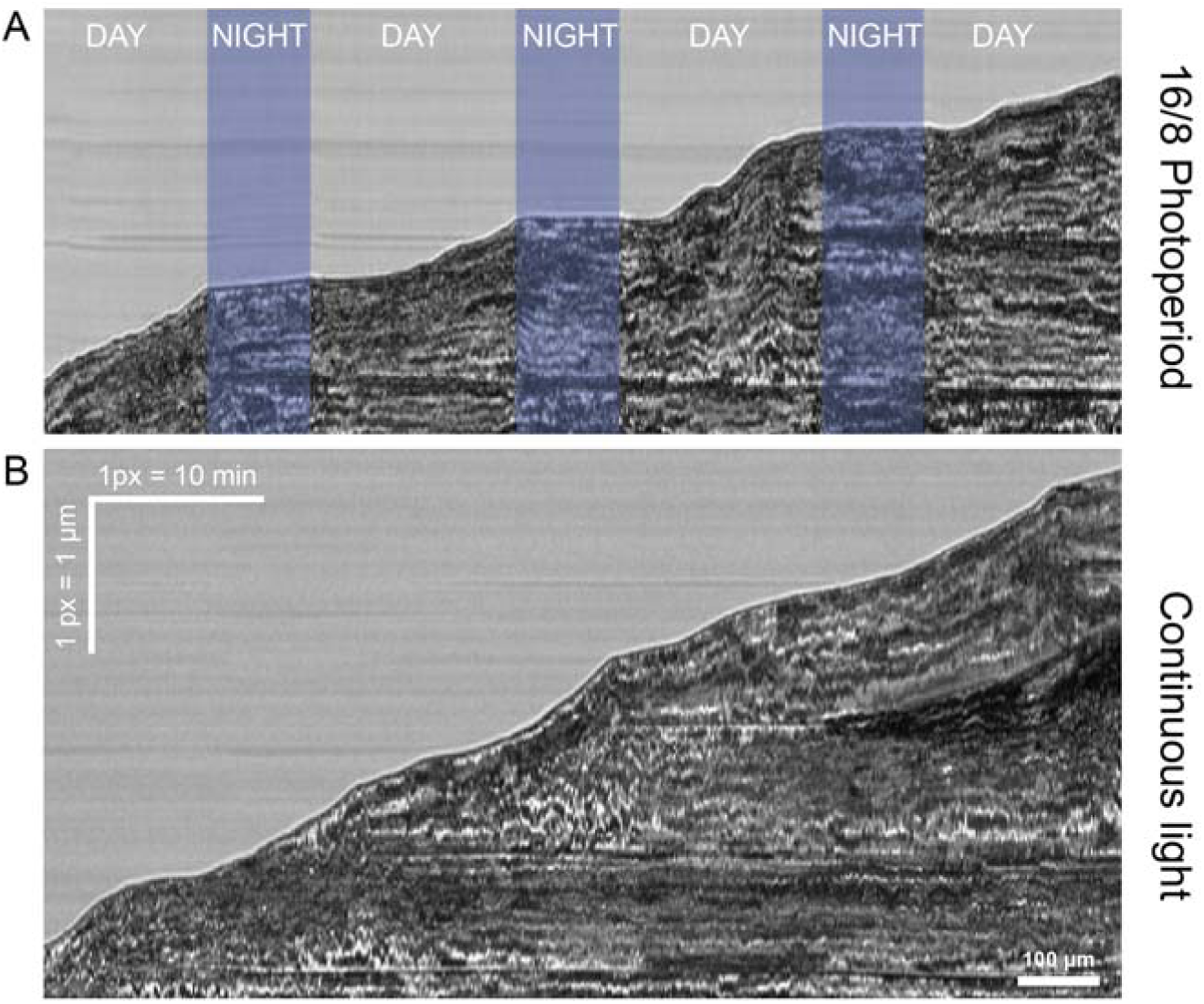
(B) Kymograph of the protonema growth under a 16-8 photoperiod. (C) Kymograph of the protonema growth under a continuous illumination. Recording have started from day 7 after protoplasts isolation their subsequent immobilization in the microfluidic traps. Protonema growth staggers during night time. Scalebars : 1 px = 10 min, 1px = 1 µm.

### Induction of leaf buds on chip using a phytohormone

As the last proof of concept of the microfluidic experimental environment presented in this work, we monitored the growth of leafy buds using the in-house microscope. Indeed, differentiation of *P. patens* caulonema to leafy buds can be induced at an early stage of development, one week after isolation and regeneration, by the addition of cytokinin (Bopp, 1984; Decker et al., 2006; Schulz et al., 2000), a phytohormone, to the culture medium. After 10 days of regeneration in the microfluidic chamber following the routine detailed in **Figure 3F**, we proceeded to a change of the PpNH4 culture medium flowing in the microchip for a 3-benzyladenin (BA) (Brun et al., 2003) in PpNH4 solution (100nM) for 96h, before finally turning back to a pure PpHN4 medium. **Figure 7** shows that the presence of BA induces the development of several buds from a filament over time. The resolution of the microscope is sufficient to observe that cell divisions occur normally in the leafy buds, and also that neighboring chlorenema stop their growth during the phyto-hormone induced differentiation, in accordance with existing literature (Bonhomme et al., 2013).

**Figure 7:**
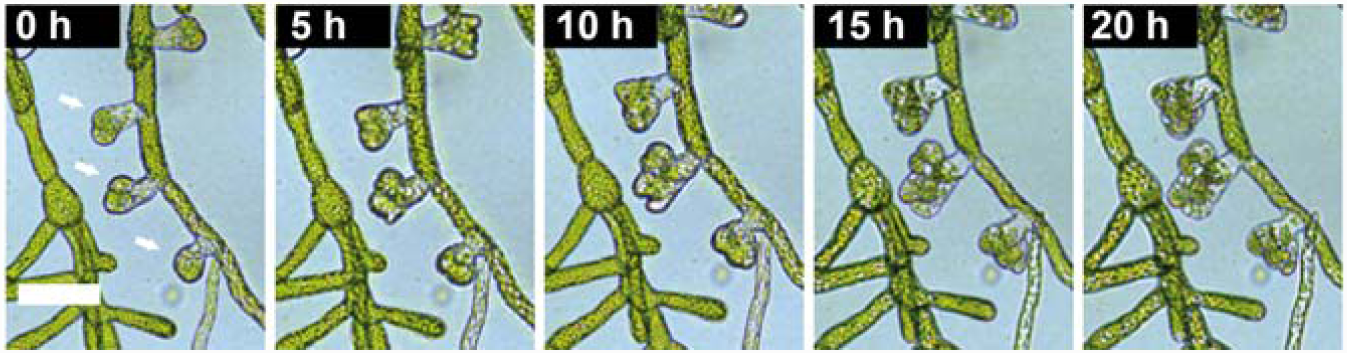
Time-lapse recording (in-house microscope) of the development of leafy buds from a *P. patens* filament after induction for 96h with PpNH4 medium supplemented with 100 nM of 3-benzyladenin. Scalebar : 70 µm.

## DISCUSSION

In this work, we present the development of a comprehensive experimental setup based on microfluidic devices made from arrays of flow-through hydrodynamic traps and allowing the long-term immobilization and regeneration of moss protoplasts and spores. Individual traps have a size comparable to the diameter of the protoplasts, and the chamber is made from a single layer of PDMS molded on a SU8-on-silicon structure, While two-layer traps can often be more efficient for trapping efficiency and homogeneity (Skelley et al., 2009), our design is more robust in terms of manipulation since after trapping, the microdevice can be moved easily from the incubators to the microscope without any risk for the developing protoplasts to move away from the traps. In addition to moss protoplasts, we have also grown spores, as their dimensions are similar. **Figure S 1** shows that spores grow nicely in the chamber, although their analysis is beyond the scope of the present study.

Instead of using syringe pumps or pressure regulators for the long-term experiments, we instead chose to use a very simple hydrostatic fluidic system that makes the samples easily transportable, straightforward to be parallelized at low cost, and less prone to contamination or sterility issues as all the elements are disposable or autoclavable. The design of the custom microscope has been driven by simplicity, so it can be very easily replicated, and avoids having cumbersome *z* defocusing over long-term observations. Its footprint is small (6×6 cm), which makes it highly parallelizable, and at the date of the report, the microscope costs less than 1000€ (ca. 1100 US$) if we exclude the prices of the Matlab licence and the computer. As Arduino boards can be easily controlled by open-source softwares such as Python, the total cost can further be reduced if needed, at the expense of a modification of the control code and the GUI.

## CONCLUSION

Herein, we developed the design of a comprehensive, physiologically relevant, easy-to-use and low-cost experimental setup for the study of *P. patens* development in a microfluidic environment. This set up allows imaging of *P. patens* development at high resolution and over long time periods. Drug or phytohormones flow-through assays can be easily achieved with the described microfluidic growth chambers and permit high-throughput pharmacological tests.

## MATERIALS AND METHODS

### Materials

Unless stated, all chemicals were purchased by Sigma-Aldrich (L’Isle d’Abeau, France) and were used as received. Antibiotics (vancomycin 50mg/L, cefotaxime 200mg/L) were used only for protonema culture on solid medium. 3-benzyladenin (CAS# 7280-81-1) was used for leafy buds induction (Brun et al., 2003).

### Microfluidic devices fabrication

For our experiments, we used two types of microchambers and trap arrays dimensions, detailed in the **Supplementary Informations**. The devices were made of PDMS (polydimethylsiloxane), using standard soft lithography techniques (Xia and Whitesides, 1998). In brief, we fabricated SU-8 masters (SU-8 2050, Microchem) on silicon wafers using a Karl-Süss MJB4 mask aligner and a laser printed photomask (Selba, Switzerland). We then proceeded to PDMS molding (1:10 ratio, RTV 615, Momentive Performance Materials) and thermal curing at 70 °C for two hours. After cutting PDMS pieces and punching out inlets and outlets with a biopsy puncher, PDMS devices were bonded to a glass bottom Petri dishes (WPI Fluorodish P35-100) using an oxygen plasma (Cute Plasma, Korea). After sealing, microfluidic chambers were washed with a solution of 0.2 wt% Pluronic F127 in water and let overnight à 4°C to make the microchannel surfaces hydrophilic.

### Plant material and tissue collection

The Gransden wild-type strain of *P. patens* (Ashton and Cove, 1977; Trouiller, 2006) and H2b-mRFP/tubulin-GFP lines (Nakaoka et al., 2012) were used in this study. Long term measurements of the influence of the light cycle on growth were done using ccd8Pp mutants (Proust et al., 2011).

### Culturing protocols

Protonemal tissue was propagated on PpNO3 medium (Ashton et al., 1979) supplemented with 2.7 mM NH_4_-tartrate. Cultures were grown in 9 cm Petri dishes on medium solidified with 0.7 wt% Agar (Kalys, France) and overlaid with a cellophane disk (Aapackaging, Preston). Cultures were grown under controlled environmental conditions: air temperature of 24.5°C with a light regime of 16 h light/8 h darkness and a quantum irradiance of 100 µE m^-2^.s^-1^ (standard conditions).

### Protoplasts isolation and purification

Isolated *P. patens* protoplast were obtained following established protocols (Schaefer and Zrÿd, 1997). In brief, protoplasts were isolated from 6-days-old protonemal cultures by incubation for 30 min in 1 wt% driselase (Sigma-Aldrich, D8037) dissolved in a 8.5 wt% mannitol solution in water. The suspension was filtered successively through 80 μm and 40 μm stainless-steel sieves. Protoplasts were sedimented by low-speed centrifugation (300 × *g* for 5 min at 20°C) and washed 3 times in a 8.5 wt% mannitol solution in water. Isolated protoplasts were then diluted to cell concentration of 2.5×10^5^ cells per mL in an iso-osmotic medium (PpNH4 medium supplemented with 6.6 wt% d-mannitol and 0.5 wt% d-glucose) prior to loading in the microfluidic device.

### Loading and regeneration in the microchip

To avoid cell and cell aggregates to stick on the sidewalls of the microtubes, we first incubate the microtubes for 10 min with a 0.2 wt% Pluronic F127 solution in water, and finally rinse them with the iso-osmotic medium described above. Then, 300 µL of fresh isolated protoplasts are pipetted in a microtube connected to the sample holder and the pressure regulator (Fluigent MFCS). Injection of the protoplasts and their subsequent immobilization took place in 3-5min at a pressure drop of 10 - 15 mBar. After loading, fresh iso-osmotic medium is flown inside the microdevice for 3-5 min, before setting up large scale reservoirs and teflon tubing used for long term culture conditions.

### Microscopy and image analysis

Long term brightfield microscopy pictures related were acquired with our custom made, in-house microscope and a 10x (n.a. = 0.10). Confocal images were acquired with a confocal Leica TCS SP8 equipped 2 Hybrid Detectors, and a 40X/1.30NA oil immersion objective. Samples were illuminated sequentially with 3 lasers at 405nm, 488nm, 552nm. Fluorescence microscope images were obtained using a Leica DMi8 microscope. Electronic Microscopy images were captured with a Hitachi TM3030 table top microscope.

### Programming, Image and data analysis

Image analysis was performed with the ImageJ/Fiji software (Schindelin et al., 2012). Data analysis was done using R (RStudio, http://www.rstudio.com/). Programming of the in-house microscope GUI was done with Mathworks Matlab and Micro-Manager (Edelstein et al., 2010) softwares.

## Supporting information

Supplementary Movie S1 - Cell wall regeneration

Supplementary Movie S2 - Protoplast division

Supplementary Movie S3 - Chloronema growth

## DECLARATIONS

### Availability of data and materials

- CIF file of the microfluidic chip mask layout
- STL design file of the 35 mm Petri dish adapter to the SM1 threading of the xy manual stage
- Matlab programming script of the microscope and Micromanager configuration file
- Connection map of the Arduino Due board
- Bill of materials of the custom-made microscope (references, manufacturers, suppliers, prices)

Documents are available on **https://github.com/FattaccioliLab/PlantsOnChip**

### Supplementary movies

- Division of a protoplast and cell wall regeneration kinetics
- Chloronemata growth under continuous illumination

### Authors’ contributions

KS, JF, JCP and FN and designed the experiments, KS performed the experiments, JF and TLS designed the in-house microscope, FC maintained *P. patens* culture, FN and SB provided the *Ppccd8* mutant. All authors participated to the analysis of the data, have read and approved the final manuscript.

## Acknowledgements

We thank B. Dubreucq (IJPB, INRA, Versailles) for the technical advices regarding the illumination setup of the in-house microscope.

## SUPPLEMENTARY INFORMATIONS

### 1. Immobilization and growth of *Physcomitrella patens* spores.

**Figure S1:**
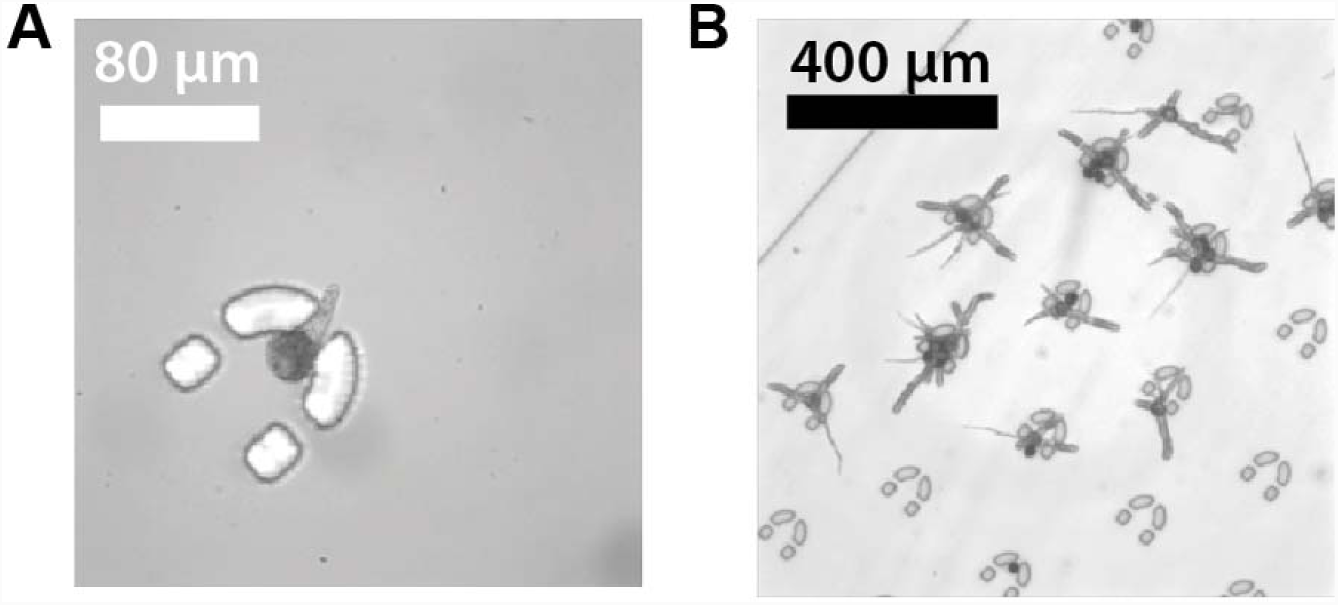
Development of P. patens spores immobilized in the microfluidic trap and fed continuously with a PpNH4 medium. (A) Germination takes places 3 days after immobilization. (D) 5 days after trapping, most of the spores have germinated and started to develop.

### 2. Effect of an overnight laser scanning confocal microscopy observation on growth

**Figure S2:**
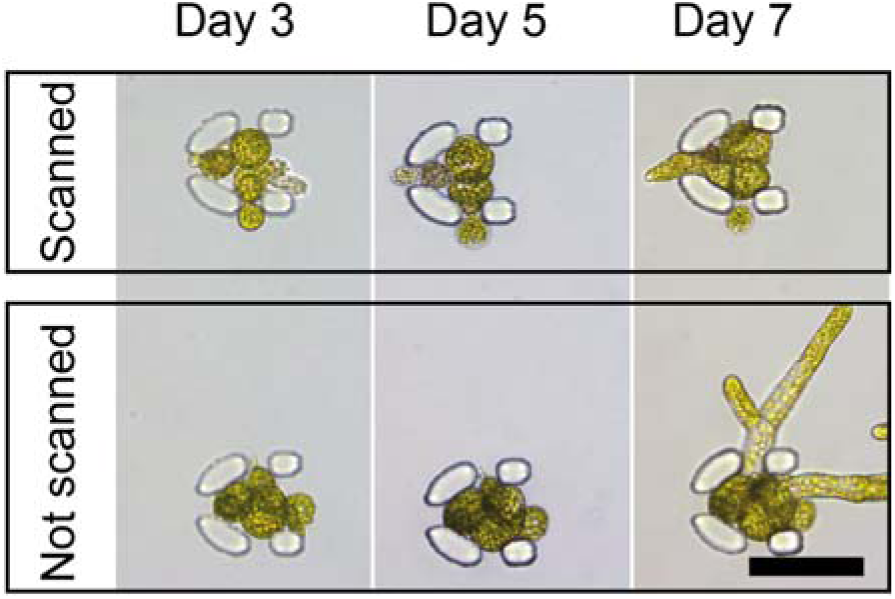
Brightfield time-lapse imaging and evolution of two adjacent traps containing protoplasts at day 3 after loading in the microchip, having (top) or not (bottom) experienced an overnight 5D laser scanning confocal microscopy session. (A) Image taken after the session, (B) 48 h after the session and (C) 96 h after the session. While the protoplasts contained in the trap that has been scanned don’t grow, the one that haven’t been scanned develop normally. Scalebar : 50 µm

